# The Latent Aging of Cells

**DOI:** 10.1101/2024.05.28.596284

**Authors:** Peter Niimi, Victoria Gould, Kyra Thrush-Evensen, Morgan E. Levine

## Abstract

As epigenetic clocks have evolved from powerful estimators of chronological aging to predictors of mortality and disease risk, it begs the question of what role DNA methylation plays in the aging process. We hypothesize that while it has the potential to serve as an informative biomarker, DNA methylation could also be a key to understanding the biology entangled between aging, (de)differentiation, and epigenetic reprogramming. Here we use an unsupervised approach to analyze time associated DNA methylation from both *in vivo* and *in vitro* samples to measure an underlying signal that ties these phenomena together. We identify a methylation pattern shared across all three, as well as a signal that tracks aging in tissues but appears refractory to reprogramming, suggesting that aging and reprogramming may not be fully mirrored processes.

## Introduction

While the underlying causes of aging are not yet understood, it presents as a progressive functional decline across systems, eventually leading to death (Kenyon 2011). A fundamental step in understanding a complex phenomenon is having metrics to characterize or quantify it. Thus, the development of biomarkers of aging is thought to be a driving force behind unlocking the potential of aging research (Marx 2024; López-Otín et al. 2013). Among the recent omics attempts to estimate the biological age of a sample/organism, DNA methylation clocks reigned supreme when compared to other modalities, such as telomere length, transcriptomic, proteomic, lipidomic, and metabolomic estimators of biological aging. DNA methylation, an epigenetic alteration associated with aging, involves the addition of a methyl group to a CpG dinucleotide (5’-C-phosphate-G-3’) (Hotchkiss, 1948; Mays-Hoopes, 1989). The associations between DNA methylation changes and aging were described as far back as 1983 in mice and a few years later in humans (Wilson and Jones 1983; Mays-Hoopes 1989; Ahuja et al. 1998); however, it wasn’t until 2011 that DNA methylation was first used to generate measures to predict aging (Bocklandt et al. 2011). By leveraging large amounts of DNA methylation data and supervised machine learning, predictors of chronological age were trained based on blood and multi-tissue DNAm samples (Hannum et al. 2013; Bocklandt et al. 2011; Horvath 2013). These made up the first generation of epigenetic clocks, which showed some initial promise as biomarkers of aging. Starting in 2018, these measures evolved from predicting chronological age to predicting more direct correlates of biological age, creating the second generation of epigenetic clocks (Levine et al. 2018; Lu et al. 2022; Hillary et al. 2020). The latest generation of clocks move one step further, swapping out the ‘ground truth’ metric upon which regression is performed, for other age-related phenomena, such as cell passaging, telomere attrition, and/or senescence (Z. Yang et al. 2016; C. J. Minteer et al. 2023; Pearce et al. 2022; Levine et al. 2019; C. Minteer et al. 2022). This begins the transition from clocks being primarily developed for the *in vivo* context into the *in vitro* realm meant to capture more mechanistic or precise components of the aging process.

Cell models have been core to biological research for decades. These models became broadly associated with the aging space following Hayflick’s proposal of the theoretical Hayflick limit (Hayflick 1965). In addition to studying the role that telomeres play in aging, cell models have been used to show and characterize cellular senescence, oxidative stress, mitochondrial dysfunction, and protein homeostasis (Bodnar et al. 1998; Gerasymchuk et al. 2022; C. J. Minteer et al. 2023; Parrinello et al. 2003). Once it was shown that methylation aging signatures tracked well across tissues and species, it was next on the docket to determine how well the measures tracked in the context of in vitro culture models. In mouse tissue, it was recently shown that serially passaging mouse embryonic fibroblasts demonstrated a similar increase in methylation as clocks applied to aging tissues (C. Minteer et al. 2022). This led to the epigenetic clock CultureAGE being generated from exhaustively passaged mouse cells. CultureAGE demonstrated surprisingly high correlation with the donor age of primary samples, despite having been trained on MEFs alone (C. Minteer et al. 2022). While an obvious explanation for the results in tissue culture could be the accumulation of senescent cells, it was shown that the measure mainly tracked mitotic rates and was not influenced by senescence induction. Further, it was later shown that epigenetic age and senescence appear to have orthogonal signatures when it comes to DNA methylation (Kabacik et al. 2022).

In addition to applying epigenetic clocks to determine factors driving aging in vitro, clocks have also been applied to examine rejuvenation effects in cultured cells. *In vitro* models show some of the strongest evidence to date in favor of the possibility of “age rejuvenation”. For instance, when adult somatic cells overexpress or are incubated with OSKM, epigenetic age predictions have been shown to drop to a near 0 age upon successful reprogramming (Gill et al. 2022). This builds on Yamanaka’s discovery that adult cells can be pushed back to an induced pluripotent state by the overexpression of Oct4, Sox2, Klf4, and c-Myc (Takahashi et al. 2007). Early epigenetic clocks showed a lot of promise in this reprogramming stage, where the epigenetic age was dramatically decreased upon reprogramming, a phenomena that was not seen with the immortalization of cells via the overexpression of telomerase (Kabacik et al. 2022). The comparison between the reset during 4F expression and the lack of reset during direct transdifferentiation of a cell lineage insinuates that aging and cell identity may have parsable features (Huh et al. 2016; Olova et al. 2019).

When considering the translation of reprogramming applications from *in vitro* to *in vivo*, one of the biggest hurdles has been the propensity for reprogrammed cells to transform. Like reprogramming and aging, cancer represents another cellular state transition that may be characterizable via methylation signatures. Literature strongly suggests that there are clear changes in methylation in cancer versus normal tissue (Gao, Widschwendter, and Teschendorff 2018; Chatterjee, Rodger, and Eccles 2018; Lakshminarasimhan and Liang 2016). However the question remains the degree to which those changes share patterns with changes observed in agingage (Lakshminarasimhan and Liang 2016; Aran and Hellman 2013; Issa 1999). To date, the rate of epigenetic change in cancer cells has been variable across different clocks, with some suggestions increasing in epigenetic age relative to the normal tissue, while others show no effect, or slight decreases (Winnefeld and Lyko 2023; Greenberg and Bourc’his 2019; Lucknuch and Praihirunkit 2022). Measuring the aging signal in abnormal tissues, such as cancers, has long proven to be a challenge with the diverse cell types at play. There is evidence that the DNA methylation signatures that track passaging *in vitro* can also differentiate tumors from normal tissues. Further, when comparing across tissues, the relative age increase in methylation signatures was shown to correlate with lifetime risk of cancer in that tissue. (Campisi 2013; C. J. Minteer et al. 2023). Overall, it has been taken to suggest that tissues prone to faster epigenetic aging may also be primed for tumorigenesis, which is further supported by the observation of accelerated epigenetic aging in normal adjacent breast tissue from women with breast cancer, relative to controls. Therefore, understanding the underlying mechanisms of age-related methylation changes in cancer presents an opportunity both in terms of biomarkers from prevention, as well as describing the phenomenological pattern of exponential increases in cancer with aging (Zabransky, Jaffee, and Weeraratna 2022).

With the goal of understanding how and why methylation changes are phenomenologically linked to aging, it will be critical to determine how these same changes link to other processes in biology, such as proliferation, stemness, and transformation. Is cancer a parallel process of aging, or perhaps is it intertwined? Relatedly, is reprogramming the complete reversal of the aging process, or do the two possess orthogonal changes? Answering such questions is essential if we want to determine to what degree reprogramming with 4F reverses aging, and whether it is constrained by de/differentiation. When we look at aging models, even cells that are seemingly immortal (i.e. by their potential to be infinitely passaged) continue to increase in epigenetic age with each passage. In light of this, it remains to be determined exactly what aspects of aging are captured by *in vitro* serial passaging (C. J. Minteer et al. 2023; Levine et al. 2019).

The aim of this paper was to computationally compare the latent DNA methylation signatures observed in aging, reprogramming, passaging, and transformation in order to assess the degree of entanglement vs. orthogonality in these phenomena. While most epigenetic clock development applies supervised machine learning in which a variable, like chronological age, is used as “ground truth” metric to train the clock, here we employ unsupervised machine learning so that results are not biased towards any specific phenotype. Our results suggest that there are methylation changes shared across aging, reprogramming (in reverse), passaging, and transformation. However, there remains a separate signature that tracks aging in tissues but appears refractory to reprogramming, suggesting that aging and reprogramming may not be mirrored processes.

## Results

A schematic for the methods used for mapping DNA methylation alterations across diverse cell types and experiments is presented in Figure 1. Training data included samples from our primary serial passaging experiment of human BJ fibroblasts (replicative senescence), and combined with public DNAm data on hRAS transformed human cells, OSKM reprogramming of human fibroblasts, and human primary sun-exposed dermal cells from young and old donors. BJ fibroblasts were serially passaged over 400 days, with samples being frozen every other passage for potential DNA methylation analysis. Senescent cells were harvested after 200 days of senescence (GSE268145). Samples were taken from the HRAS cohort to model H-ras-V12 oncogene induced senescence, xenografted cells, and sV40 combined with hTert immortalization (GSE91069). Reprogramming through OSKM overexpression was induced in primary skin samples via retroviral transduction, and TRA-1-60+ cells were profiled for DNAm on days 7, 11, 15, 20, and 28 (GSE54848). Samples from young (ages 20-34) and old (ages 65-90) donors were collected for paired sun-exposed and sun-protected dermis and epidermis and subjected to DNAm profiling (GSE52980).

**Figure 1:**
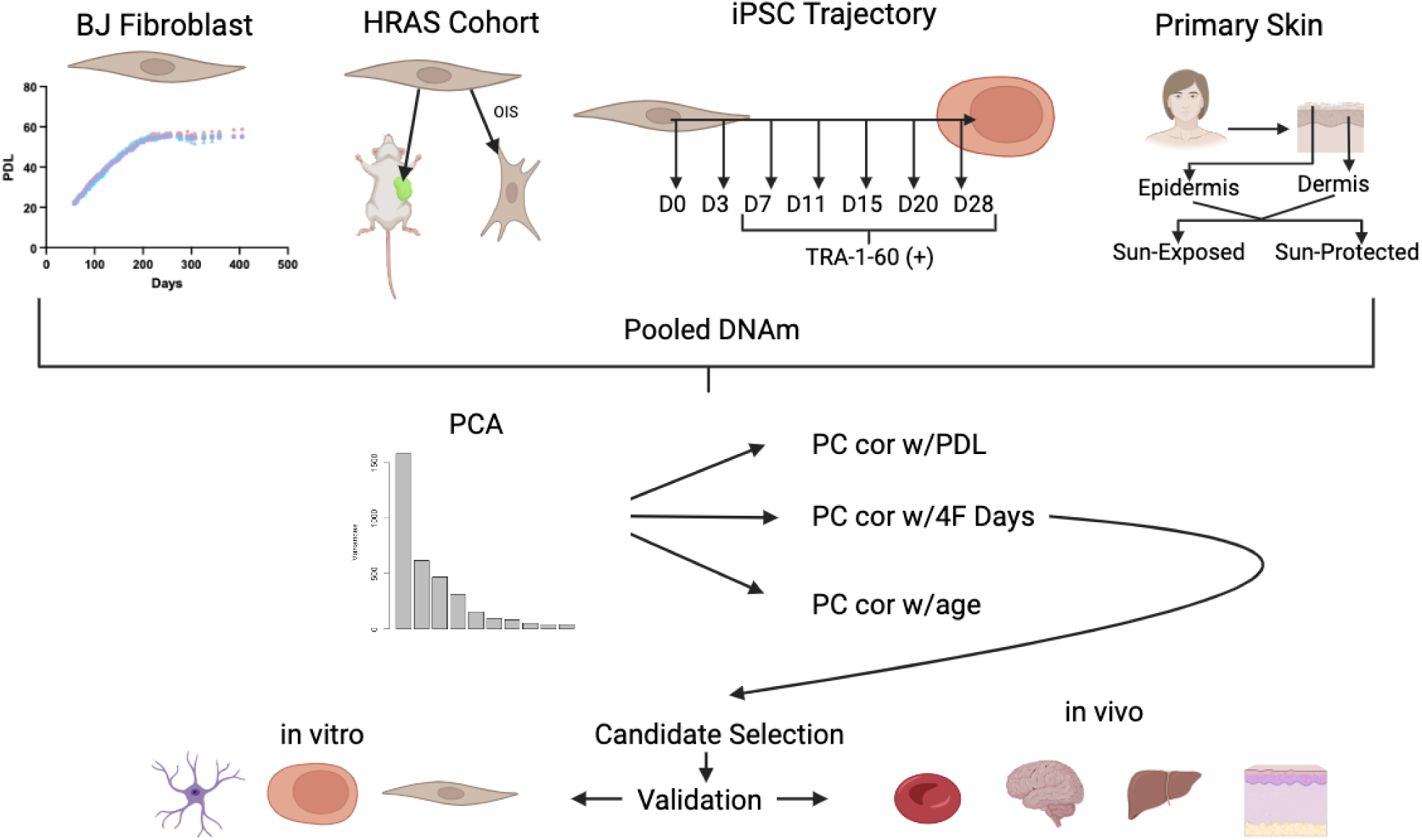
Project Workflow Schematic for data collection through data processing. Created with BioRender.com.

For our training data, we pooled methylation data generated from a subset of the BJ fibroblast serial passaging (consisting of n=10 serially passaged BJ fibroblast samples), n=17 samples from an HRAS cohort, a subset of n=18 iPSC time course samples, and n=20 sun exposed dermis samples. The BJ fibroblast title refers exclusively to the serially passaged BJ fibroblasts and not BJ fibroblast cell line derivatives that are found in public datasets such as the HRAS cohort. Principal component analysis (PCA) was then performed on the pooled DNAm data to identify latent signals capturing variance across these diverse samples. This produced latent variables, hereby referred to as ELDAR (Epigenetic Latent signals of Differentiation, Aging, and Reprogramming).

The PCs from ELDAR were then correlated with each of the time dimensions in the independent datasets (except for the HRAS cohort which lacked time information) to assess the degree they captured variance in aging, replication, and reprogramming. Specifically, the time dimensions reflect: the age of the donor for the primary dermal samples; population doubling level for BJ fibroblasts; and days after Oct4, Sox2, Klf4, c-Myc (4F) induction for the iPSC trajectory, respectively.

### Time Correlations in ELDAR

As shown in figure 2A, PCs 1-4 tend to show strong time correlations for skin aging and/or reprogramming (greater than abs(0.5) on the x-axis and/or the y-axis). For PCs 2-4, high time correlations are mostly observed for the reprogramming axis, while little to no correlation is seen for aging in skin. However, for PC1 we observe a strong inverse relationship between the correlations, suggesting that this signal increases with aging (r = 0.73) and decreases with reprogramming (r = -0.93). PC1 correlations also show positive associations in BJ passaging and skin aging (Figure 2B), as well as inverse correlations in reprogramming and passaging (Figure 2C). Again, this suggests that PC1 increases with aging and passaging and decreases upon reprogramming. Conversely, PC5 shows moderate correlation with aging and passaging in the shared direction and only a moderate correlation with reprogramming in the inverse direction (Figures 2A-2C). For PCs 2-4, correlations across the three domains (aging, reprogramming, and passaging) are either weak or inconsistent.

**Figure 2:**
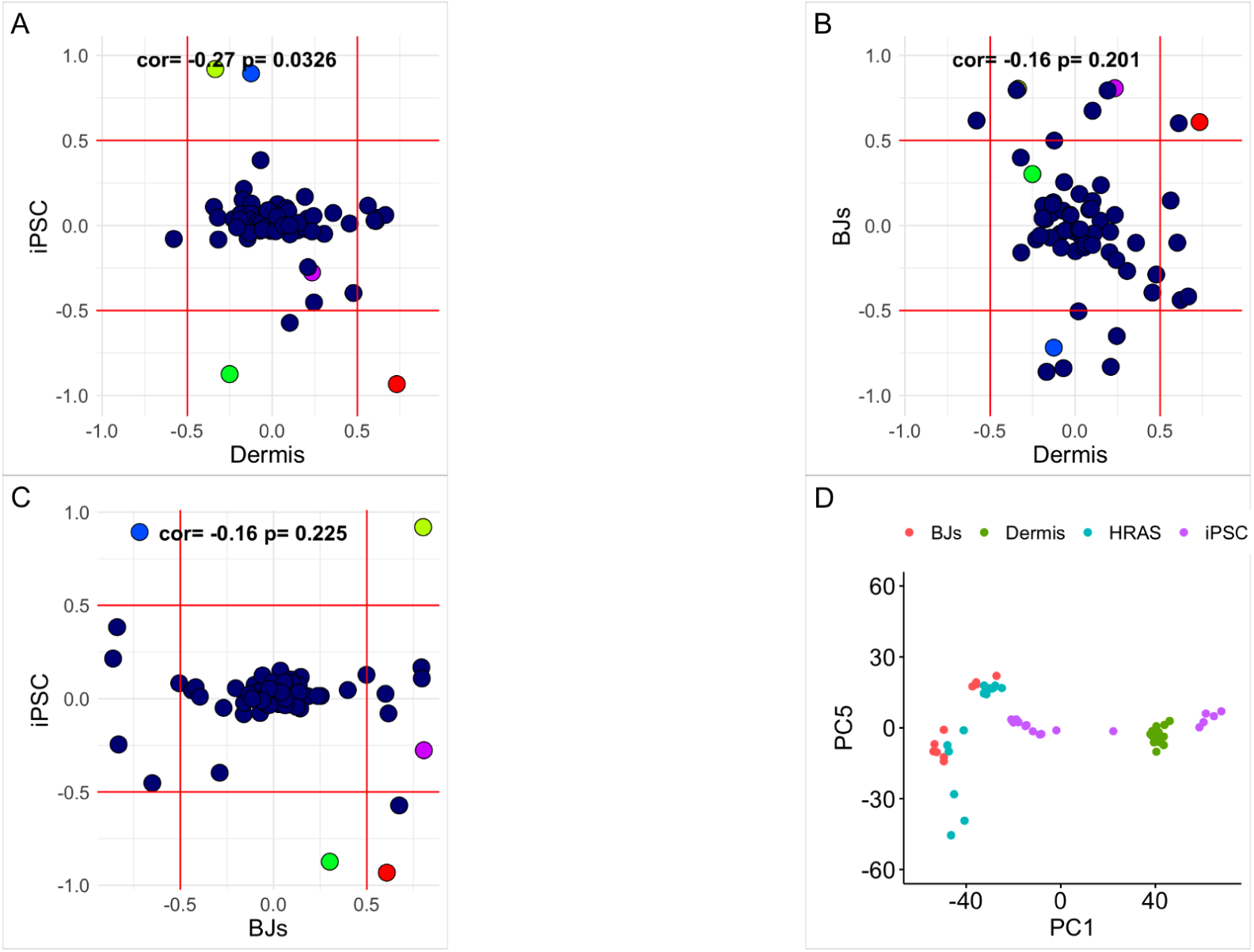
Selection of PCs PC Time correlations for iPSC v Dermis, BJs vs Dermis, and iPSC vs BJs. The first 5 PCs in the correlation plots are red, yellow, green, blue, purple respectively. The last plot is the PC1 vs PC5 variance plot for serial passaged BJ fibroblasts in red, sun-exposed dermis in green, HRAS cohort in blue, and iPSC time course in purple.

Figure 2D shows various training samples plotted as a function of PC1 and PC5. These results highlight that reprogramming cells exhibit a large variance in the PC1 dimension, but almost no variance across PC5. Conversely, skin aging samples tend to have slightly higher variance across PC5 compared to PC1. Passaging cells and HRAS transformed cells (vs. controls) exhibit variability across both PC1 and PC5 dimensions. Taken together, this suggests that the signals captured by PC1 and PC5 may change in both *in vivo* and *in vitro* models of aging, but only those captured by PC1 exhibit variance upon reprogramming.

### ELDAR in Primary Tissues

PC1 and PC5 from the initial dataset were then projected onto independent validation data to assess changes with aging and/or rejuvenation across tissues and cell lines. For data from tissues, these scores were each plotted against the age of a sample. The unsupervised measures captured by PC1 had moderate to weak, but consistent correlations with age in the liver, brain, blood, dermis, and epidermis (Figure 3.1). Of these tissues, the PC1 score had a correlation with tissues that tend to be prone to damage like the skin, and also correlated with the developing brain. Intriguingly, ELDAR was very sensitive to the damage relating to skin’s sun exposure, exhibiting a higher rate of increasing PC1 scores as a function of aging (p =1.20e-08).

**Figure 3:**
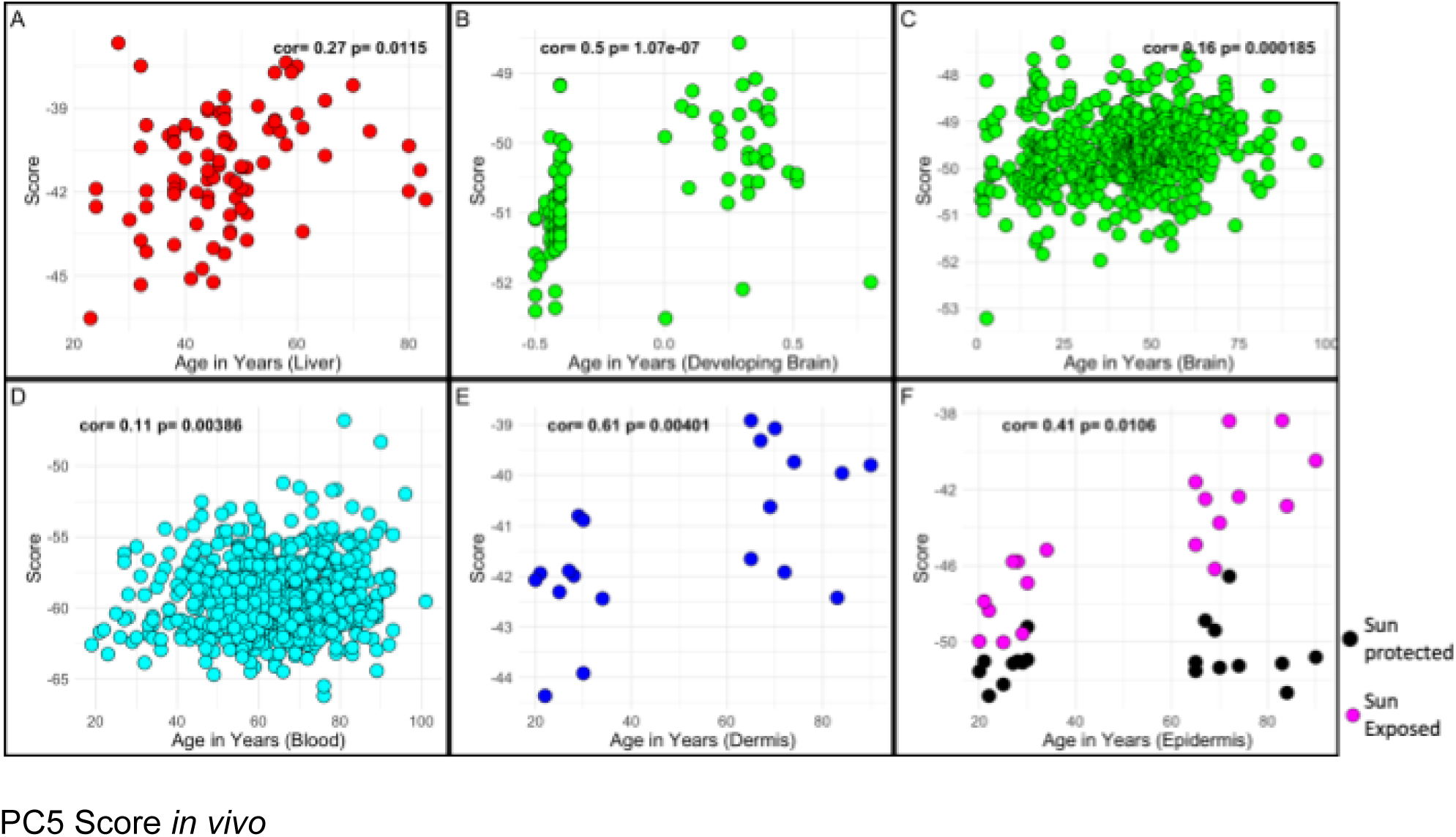
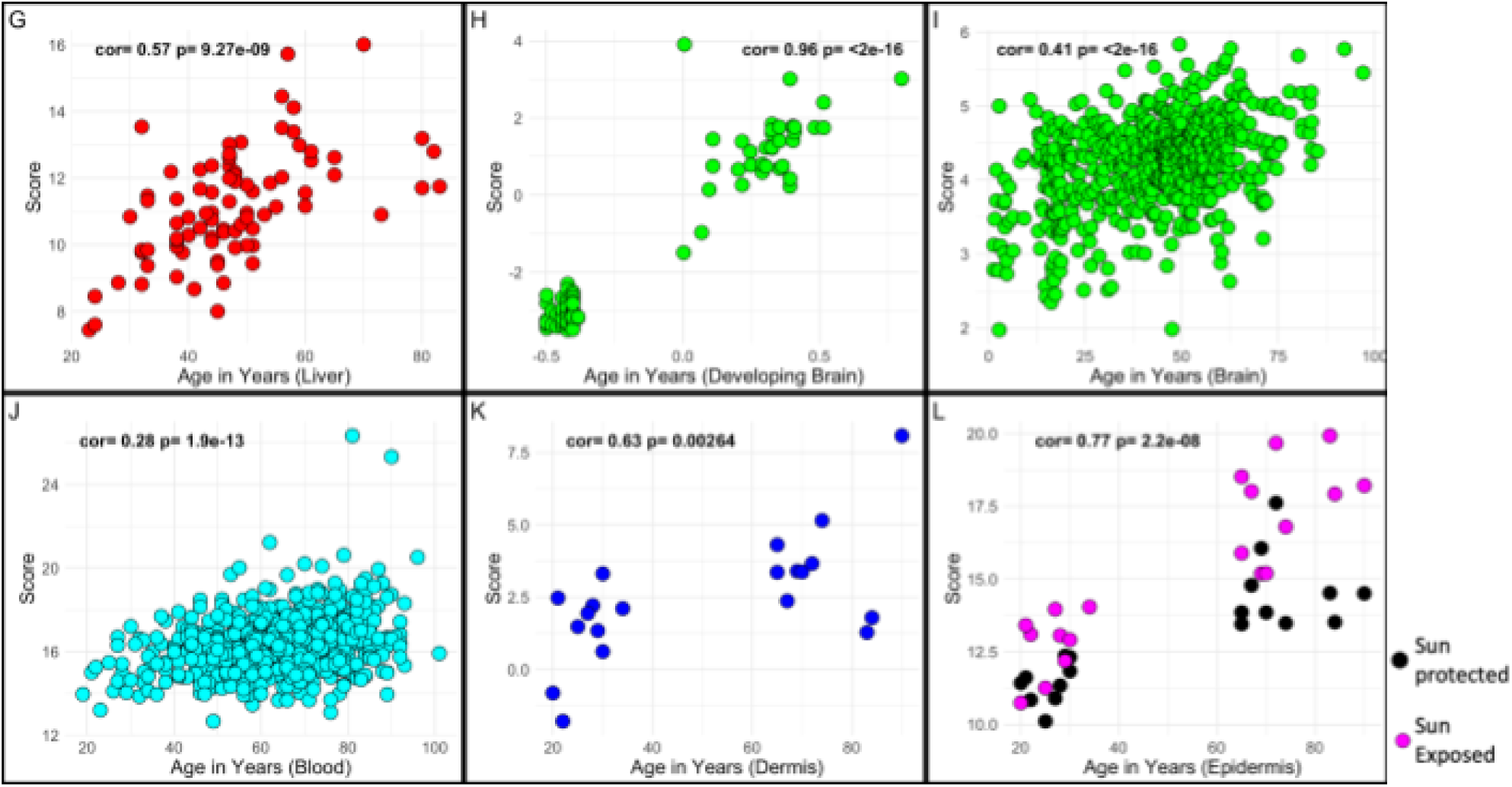
PC1 Score *in vivo* Liver samples shown in red (GSE48325), brain samples shown in green (GSE74193) separated by samples under the age of one as developing, blood samples in cyan (GSE40279), sun-protected dermis samples in blue (GSE52980), sun-protected epidermis samples in black (GSE52980), and sun-exposed epidermis samples in pink (GSE52980).

The PC5 score had a higher correlation with age than the PC1 score in all the tissues except the dermis (Figure 3.2), with the strongest age correlation found for the brain during development (r=0.96), despite having no developmental samples and no brain samples in the training data. Similar to PC1, we also observed an effect of sun exposure in epidermis, in which sun-exposed samples exhibited increases in PC5 overall (p=5.42e-10), and also an accelerated increase in PC1 as a function of age and sun-exposure (p=3.99e-5).

### ELDAR in Reprogramming

Next, we examined the patterns of PC1 and PC5 scores during OSKM reprogramming. There exists speculation that reprogramming constitutes aging reversal, suggesting that it fully reverses the processes of aging (J.-H. Yang et al. 2023). If true, one would anticipate to observe negative correlations as a function of reprogramming time, suggesting that the age score is reset to a more youthful state. The PC1 score showed an expected reduction during lentiviral infection induced full reprogramming with OSKM (p=1.79e-08), partial reprogramming with transient dox-inducible OSKM (p=0.000433), and Sendai virus mediated OSKM treatment (p=0.00422). The PC1 measure also differentiated the cells that were successfully reprogrammed by the Sendai viral cocktail from those that were classified as “failed to reprogram”.

If this resetting was a universal signature of reprogramming, then one would anticipate that all age correlated PCs would show a robust negative correlation with reprogramming time. Yet, in PC5, reprogramming does not induce a reduction in the score inverse of the association with aging. In fact, our results suggest that in the initiation phase of reprogramming–commonly associated with partial (or transient) reprogramming– the cells appear to be increasing in PC5, indicative of an increased aging phenotype. This is observed both in the early-mid phases of the lentiviral treated cells and in the dox-inducible OSKM transiently reprogrammed cells (Trans4F - GSE165180). However, this effect appears to revert in the lentiviral treated cells, beginning during the maturation phase, which is commonly associated with dedifferentiation (Buganim, Faddah, and Jaenisch 2013). The PC5 score was also unable to differentiate between the successful and failed to reprogram Sendai conditions. This together points to an underlying feature of aging that is captured by DNA methylation, yet is not amenable to restoration via existing epigenetic reprogramming methods, at least prior to the loss of cell identity.

**Figure 4:**
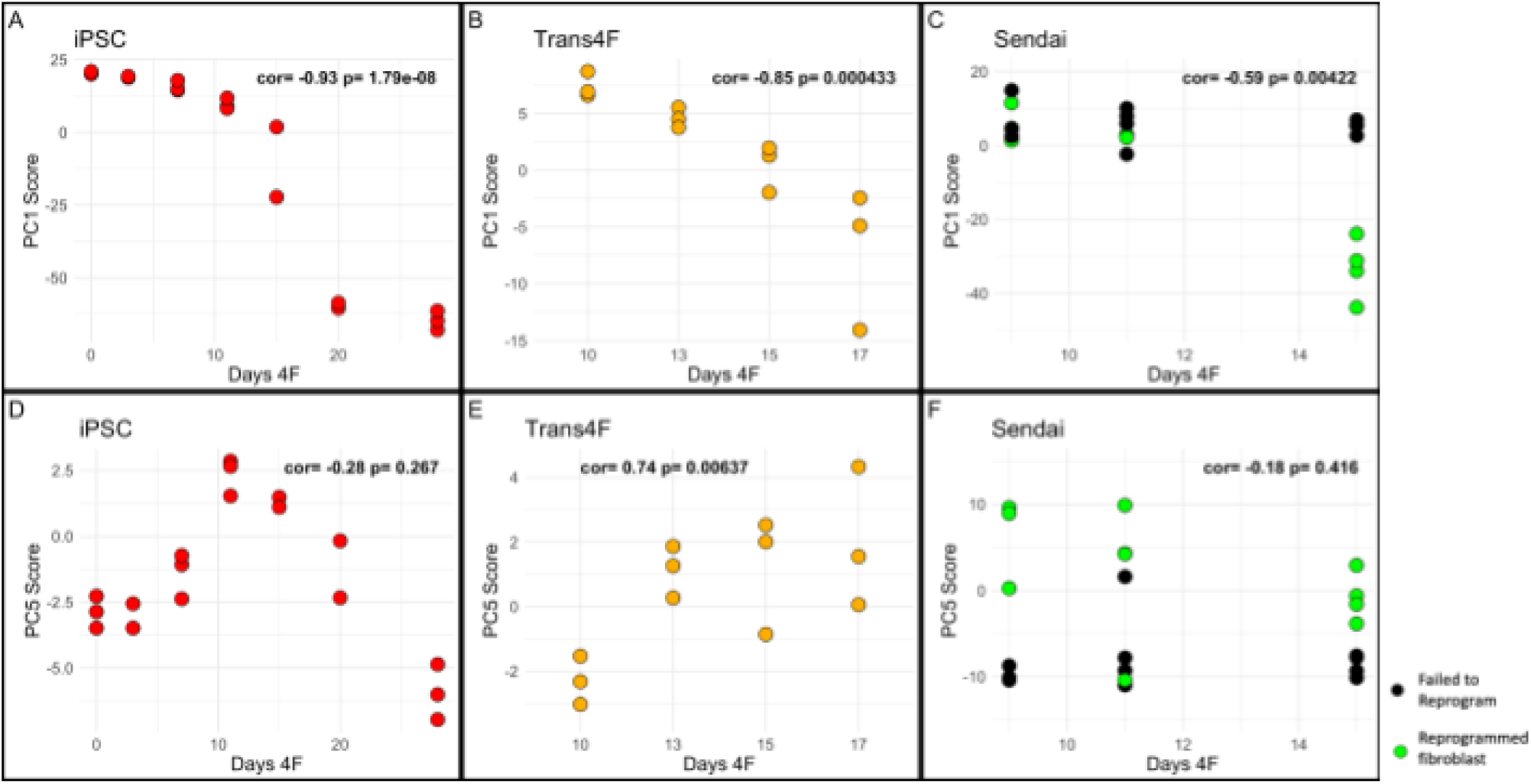
PC1 and PC5 plotted against days of OSKM expression in 3 models of reprogramming iPSC time course in red (GSE54848), transient reprogramming by dox-inducible OSKM time course in yellow (GSE165180), Sendai time course failed to reprogram in black (GSE165180), and Sendai time course successfully reprogrammed in green (GSE165180).

### ELDAR in Cultured Samples

To ensure that the divergence in behavior of the reprogramming samples was not simply a result of the difference between cultured cells and primary samples, the PC1 and PC5 scores were applied to a variety of cultured samples. The first sample (GSE158089) started as iPSCs that were then differentiated into neural progenitor cells (NPC) at day 16 and then into neurons at day 37 and then maintained through day 58 as neurons. PC1 showed a strong significant correlation with an increase in days in culture, indicative of the differentiation process (r=0.78, p=1.1e-3). For PC5, a trend was observed in the differentiating neuronal sample (r=0.42), however, it was not significant (p=1.4e-1). PC1 and PC5 were examined as a function of cPD in both cultured BJ fibroblasts and fetal astrocytes (GSE226079). The BJ Fibroblasts included those used in training (set 1), as well as those derived from an independent passaging experiment using a different donor line (set 2). For all serial passaging samples, we observed strong correlations for both PC1 and PC5 with cPD levels ranging from r=0.72 (PC1, BJ set 1) to r=0.95 (PC1, BJ set 2 and PC5, astrocytes) with population doubling levels. This trend also persisted with the expression of hTert, which was used to immortalize both the astrocyte (PC1: r=0.88, p=3.08e-15; PC5: r=0.91, p<2.00e-16) and BJ fibroblasts (PC1: r=0.92, p=1.26e-9; PC5: r=0.95, p=2.08e-11). Interestingly, in differentiated neurons, astrocytes, and the BJ fibroblast set 1, we observed a trend of the score leveling off once the population of cells stops dividing, either through differentiation into postmitotic cells or via the onset of cellular senescence.

**Figure 5:**
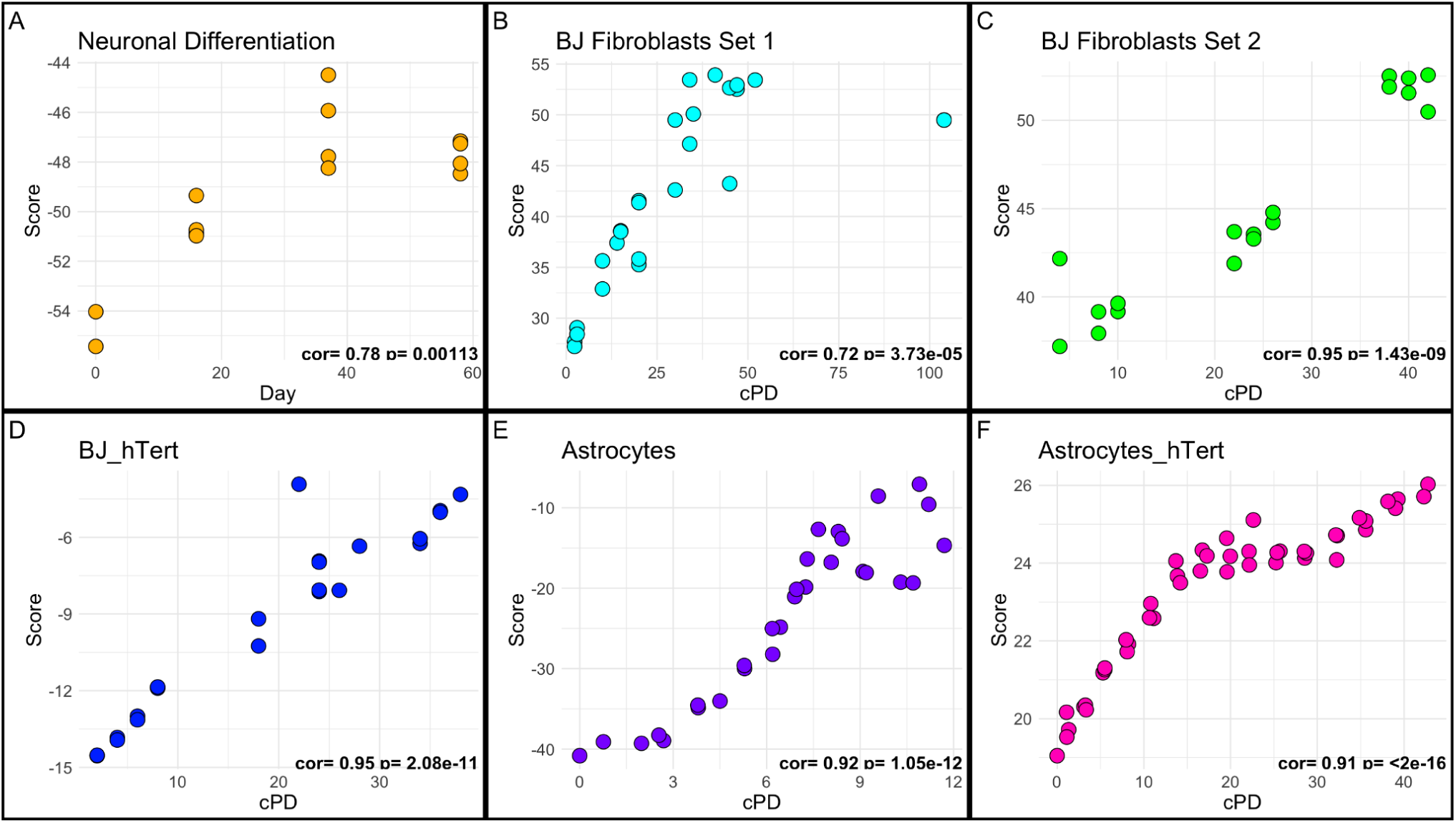
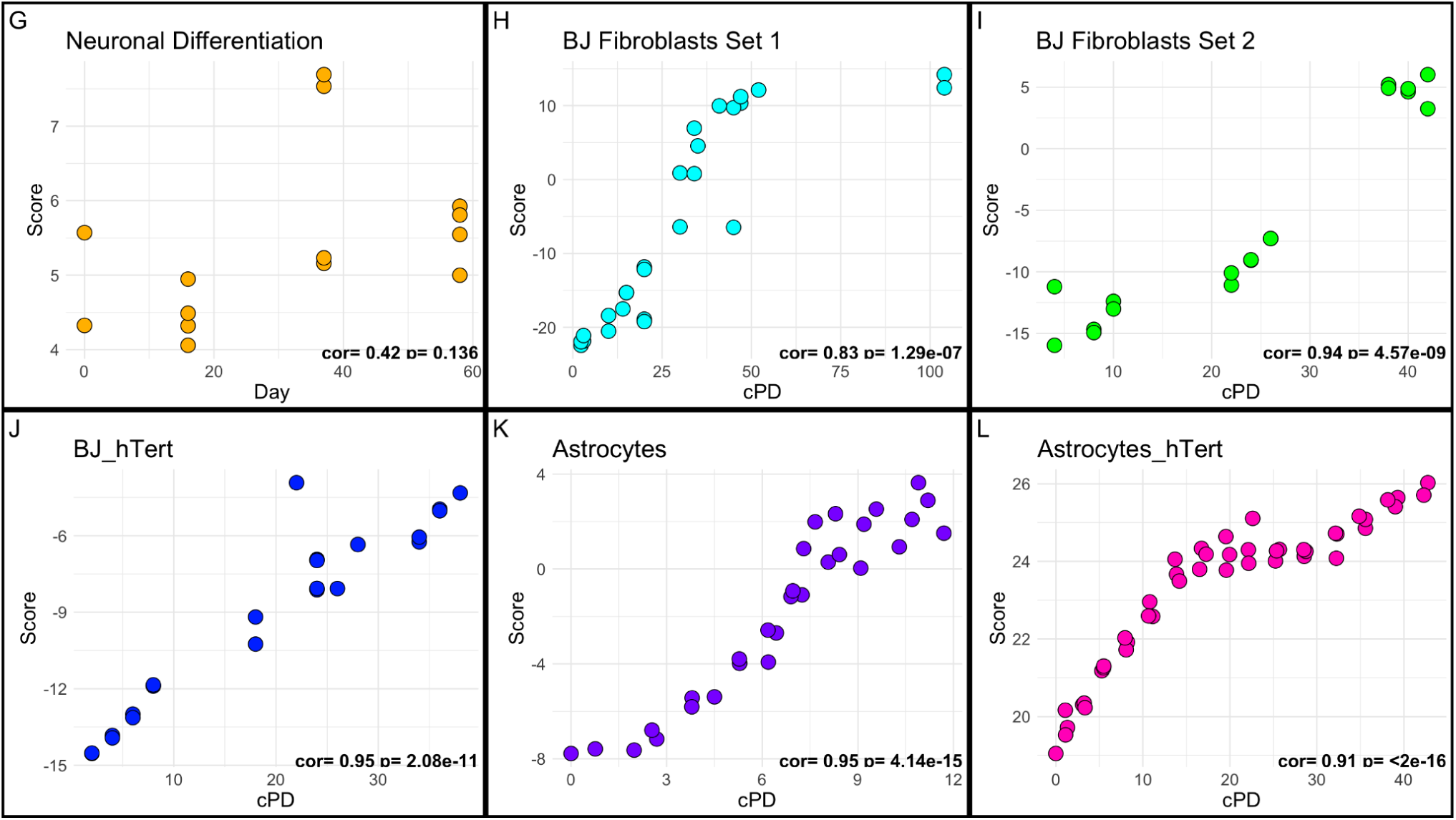
PC Scores *in vitro* iPSC differentiation into neuronal cell line in yellow (GSE158089), BJ fibroblast serial passaging set 1 in cyan, BJ fibroblast serial passaging set 2 in green, serially passaged hTert immortalized BJ fibroblasts in blue, serially passaged astrocytes in purple (GSE226079), and serially passaged hTert immortalized astrocytes in pink (GSE226079). PC1 scores shown in A-F, PC5 scores shown in G-L.

### ELDAR CpG’s Chromatin State & Location Enrichment

To identify regions and chromatin states driving these candidate aging signals, the top 1000 CpGs each with the highest positive and negative loadings from both PC1 and PC5 were selected. The CpGs with positive loadings reflect those that tend to gain DNAm with aging and in the case of PC1, decrease DNAm with OSKM treatment. Alternatively, the negative loading CpGs lose DNAm with aging and, for PC1, regain DNAm with reprogramming. ChromHMM was used as a reference to determine enrichment of chromatin states within the four groups (positive and negative loading CpGs, from PC1 and PC5 separately). The locations of the CpGs are defined as; CpG islands: regions with a high frequency of CpGs, often associated with gene silencing, CpG shores: regions up to 2kb from a CpG island, CpG shelves: regions from 2kb to 4kb away from CpG islands, and CpG open sea: CpG sites that are relatively isolated from CpG islands. The percentage of each location type was then normalized to the background of all the CpGs considered in our initial analysis. Results suggest that bivalent promoter and enhancer regions were the most enriched among positive loading CpGs (i.e. those that gain DNAm with aging) for both PC1 and PC5 (PC1 enrichment=4.52-fold; PC5 enrichment =2.26-fold). These CpGs were composed of mostly CpG islands. However, those in PC1 also exhibited a high fraction of CpGs in Shores.

For negative loading CpGs (those that decrease with aging), we find that PC5 is enriched for Open Seas in active enhancer regions (enrichment =3.46-fold, as well as in acetylated regions (enrichment =2.46-fold). PC1 loss of DNAm aging CpGs on the other hand, showed greatest enrichment for quiescent state CpGs (enrichment =8.6-fold), GapArtf which denote those with minimal chromatin marks (enrichment =3.53-fold), weak enhancers (enrichment =3.17-fold), active enhancers (enrichment =2.68-fold), and regions of acetylation (enrichment =2.74-fold). The two PCs also differed somewhat in the locations of CpGs with high or low loadings. For instance, CpGs with age-related gain in PC1 were found primarily in Islands (48.4%) and Shores (41.3%). Conversely, those in PC5 were in Open Seas (45%), although with minimal enrichment beyond background frequency. However, for both PCs the CpGs that tend towards loss of DNAm with aging were enriched in Open Sea regions, which tend to be CpG sparse (PC1=82.4%, PC5=71.8%).

**Figure 6:**
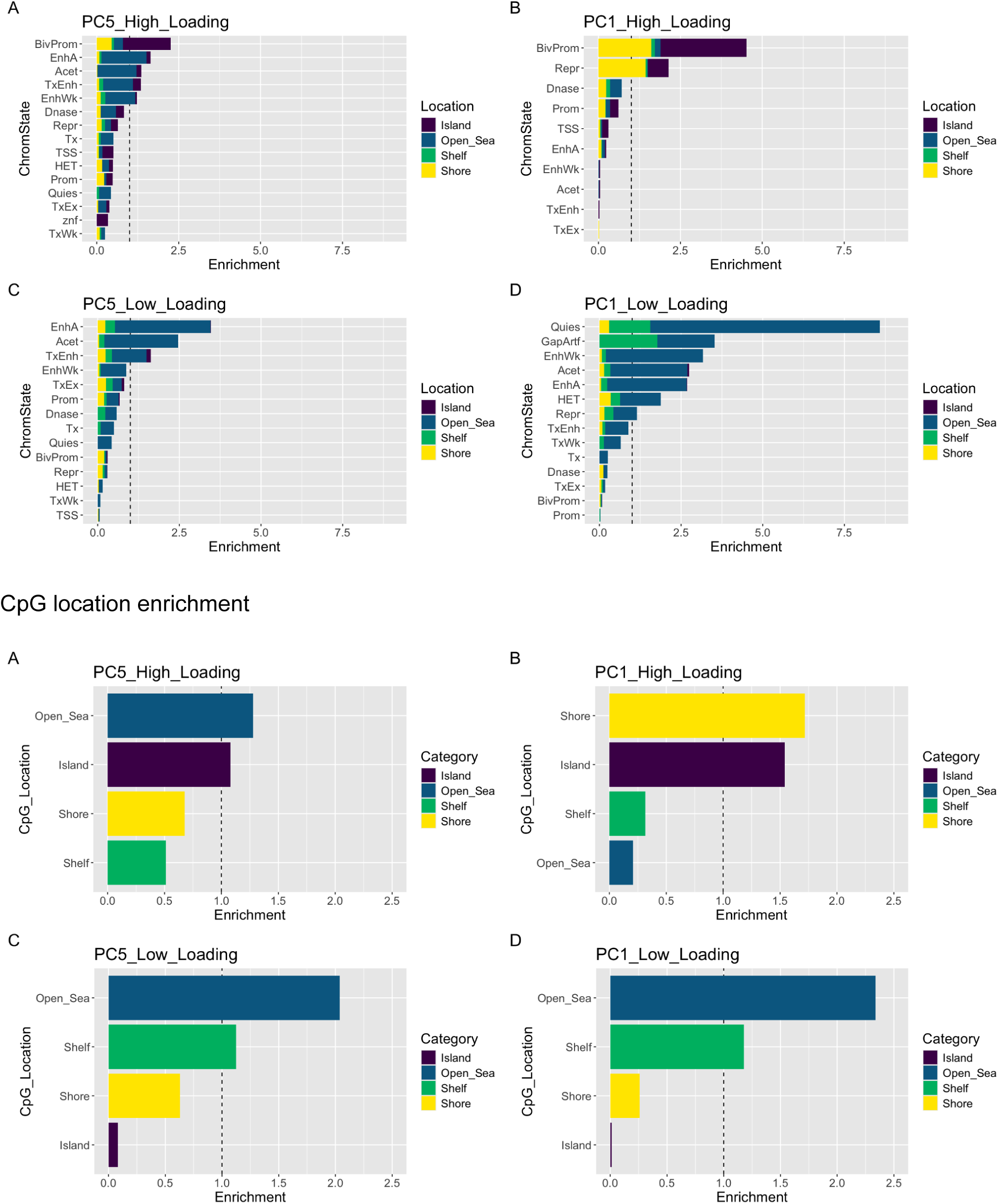
Chromatin State Enrichment CpG location enrichment for the top and bottom 1000 loading CpGs in PC1 and PC5 respectively. CpG islands in purple, open sea CpGs in blue, shelf CpGs in green, and shore CpGs in yellow.

### Comparison of ELDAR to Existing Clocks

Finally, we compared the overlap in the top CpGs from ELDAR to sets of CpGs found in existing epigenetic clocks. From the 3947 CpGs that made up the top and bottom 1000 loading CpGs in PC1 and PC5 there was no overlap with the Horvath Pan-Tissue clock, a 3.8% overlap with the Horvath Skin-Blood clock, a 5.6% overlap with the Hannum clock, and a 0.8% overlap with PhenoAge (Table 1) (Hannum et al. 2013; Levine et al. 2018; Horvath 2013; Horvath et al. 2018). Whereas the overlap with the 4 clocks to each other ranged from 1.1% between the Hannum and PhenoAge clocks to 15% between the Horvath Skin-Blood and PhenoAge clocks (Table 2). Given that the selection of the 1000 highest and lowest CpGs could bias inclusion of CpGs, we additionally investigated the relative density where all the clock CpGs fall within the loadings distributions for PC1 and PC5 (Supplemental Figure 1). Similar to the percentages, the results suggest the Hannum and the Horvath Skin-Blood clocks were more likely to contain CpGs with higher absolute loading values, while the Horvath Pan-Tissue and PhenoAge clock CpGs tended to have lower loadings.

**Table 1:** PC1 associations with age and sun exposure.

|  | Coefficient | P-Value |
| --- | --- | --- |
| Age | 0.12009 | 1.2e-8 |
| Sun Exposed | -0.54425 | 0.676 |
| Age * Sun Exposed | -0.10714 | 3.99e-5 |

**Table 2:**
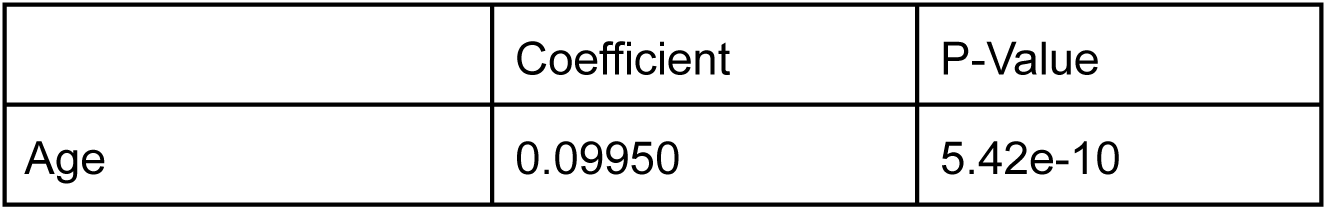
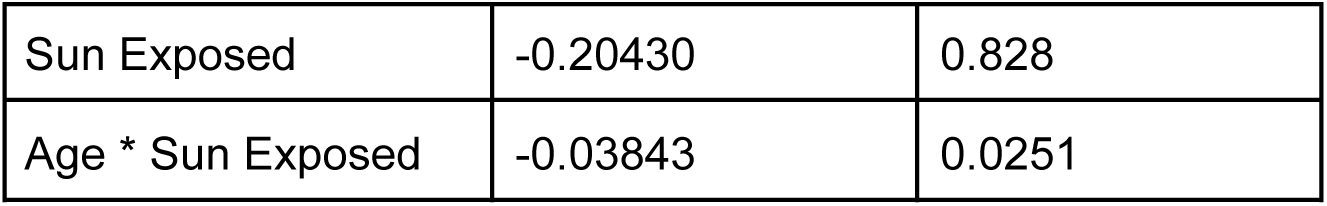
PC5 associations with age and sun exposure.

**Table 3:** ELDAR’s overlap with the Horvath Pan-Tissue, Horvath Skin-Blood, Hannum, and PhenoAge clocks.

| CpG Association | Horvath Pan-Tissue | Horvath Skin-Blood | Hannum | PhenoAge |
| --- | --- | --- | --- | --- |
| Age-Increasing PC1 | 0 | 6 | 1 | 1 |
| Age-Increasing PC5 | 0 | 4 | 3 | 0 |
| Age-Decreasing PC1 | 0 | 0 | 0 | 1 |
| Age-Decreasing PC5 | 0 | 5 | 0 | 2 |

**Table 4:** Overlap between the Horvath Pan-Tissue, Horvath Skin-Blood, Hannum, and PhenoAge clocks.

| Epigenetic Clocks | Horvath Pan-Tissue | Horvath Skin-Blood | Hannum | PhenoAge |
| --- | --- | --- | --- | --- |
| Horvath Pan-Tissue | 353 | 60 | 6 | 41 |
| Horvath Skin-Blood | 60 | 391 | 45 | 58 |
| Hannum | 6 | 45 | 71 | 6 |
| PhenoAge | 41 | 58 | 6 | 513 |

## Discussion

Since the first development of epigenetic clocks in 2011, the field has applied the measures to study a wide array of questions spanning from aging,disease risks, treatments, and even social factors (Bocklandt 2011,Marx 2024). To date these clocks have evolved to be targeted for predicting a diversity of outcomes including chronological age, biological age, morbidity/mortality, and cellular hallmarks based on a training regression to specific variables or clinical measures (Bergsma and Rogaeva 2020; Lu et al. 2022; Levine et al. 2018). While this may produce specialized measures that allow for precise predictions of the phenotypes they were trained on, they may fail to capture other meaningful aspects of the multi-factorial biological aging process. Most, if not all, epigenetic clock development to-date has applied supervised machine learning in which a variable, like chronological age, is used as “ground truth” to train the clock. However, this may make clocks less valid or biased when testing for the intersection of aging signals and other phenomena, like reprogramming, cancer, differentiation, or replication.

Here we employ unsupervised machine learning to generate an unbiased measure that isn’t overly fit to any one specific phenotype, and instead, enables us to identify methylation changes shared across aging, reprogramming, passaging, and transformation. In doing so, we were also able to separate signatures that track aging in tissues but differ in their modulability by reprogramming, suggesting that aging and reprogramming may not be perfectly mirrored processes. Our findings suggest that the CpGs captured in PC5 that are altered in aging and not reset with reprogramming may be part of the epigenetic ‘memory’ that others have previously described when comparing iPSCs to ESCs(Plath 2011,Reik 2014). While once thought to be a failing of the reprogramming process, this could be one of reprogramming’s inherent safeguards (Abad et al. 2013). This implies that no matter how much work is done to optimize OSKM reprogramming efficiency, one may never get a full population recovery back to the basal state without the application of new pioneer transcription factors. While beyond the scope of the current study, it is possible that other protocols using novel factors could be discovered that are able to reset the aging changes captured in PC5 given that one must assume these aging signatures are absent in the ground state of the developing embryo (Gladyshev 2021). Alternatively, it is possible that gametes may be resistant to these age changes or that selection can differentiate aged vs. youthful cell states and thus nature would not have needed to optimize a method to reset them.

Regardless, given the lack of mechanistic understanding for epigenetic clocks, the relative importance of CpG signatures that are not responsive to OSKM reprogramming versus those that are remains unknown. For instance, it may be the case that such CpGs are not causally implicated in aging phenotypes and therefore, the inability to reset them is inconsequential. Further, if they are meaningful for aging biology, our inability to specifically target them may continue to limit the utility of reprogramming interventions. Our group has previously reported evidence of this divergence in the behavior of DNA methylation during reprogramming (Levine 2022). We demonstrated that CpGs used in epigenetic clocks can be broken down into subsets of 12 unique modules, each of which favors a biological readout. These results culminate with the identification of a subset of underlying aging signatures that current reprogramming procedures are unable to target. However, we also showed that these modules were less predictive of remaining lifespan compared to those that were responsive to OSKM treatment.

Another outstanding question in regards to reprogramming as an anti-aging intervention centers around the entanglement between differentiation and aging. While cells are thought to be rejuvenating in response to OSKM treatment, they are also undergoing dedifferentiation into a pluripotent state. Yet, it remains unknown whether the changes observed across the reprogramming time-course reflect one or another of these processes or even both. Further, it is unknown whether there is a limit to the degree of rejuvenation one can achieve without dedifferentiation–at least transiently (Gill et al. 2022). As such, we tested the degree to which our signatures were also responsive to differentiation. Using an iPSC derived neuronal differentiation model that takes 60 days to mature, we saw that PC1 remained correlated with the process over time while PC5 was less so. When combined with the resetting of the PC1 signature in OSKM reprogramming, PC1 may be more linked to dedifferentiation than PC5. We also observed insignificant changes between the day 37 and day 58 neurons in either PC, which is in agreement with the authors’ original conclusions that showed a negligible difference between the neuronal samples and the progenitors (Imm et al. 2021). Finally, we observed a potential plateau in both PC1 and PC5 upon terminal differentiation. In moving forward, these observations should be confirmed in a dataset that continues to sample terminally differentiated cells in culture over longer periods of time.

The observation that both PCs plateau once cell division ceases was also consistent with our observations in serially passaged cells. For instance, in both fibroblasts and astrocytes, we observed increases in PC1 and PC5 in association with population doubling. However, our results suggested that once cells entered senescence PC1 and PC5 again plateaued. Further, hTERT immortalized cells exhibited increases that continued for the duration of the passaging experiments. This is consistent with previous results from epigenetic clocks demonstrating that DNAmAge increases as a function of mitotic rate, yet the signal is not accelerated by the induction of cellular senescence (C. Minteer et al. 2022; Kabacik et al. 2022). This pattern observed both here and in epigenetic clocks may reflect DNAm changes associated with hypermethylation of polycomb-associated promoters.

In biology, the gain of DNA methylation in promoters is associated with a silencing of gene expression while the loss of DNA methylation in enhancers allows for transcription factors to bind and induce downstream gene expression (Lakshminarasimhan and Liang 2016). In our data we show that the regions which are gaining the most methylation are the bivalent promoters that are primarily found in CpG islands. This is in agreement with prior reports in normal aging (Johnson et al. 2012; Rakyan et al. 2010). It’s also notable that these changes have been associated with cancer development, and states more common to iPSCs than to wild type cells (Bernhart et al. 2016).

Despite the similar patterns in CpG characteristics between ELDAR and existing epigenetic clocks, we observed very few shared CpGs. This is consistent with prior work describing distinct CpG sets across clocks (Duan et al. 2022). Yet,despite having little overlap with one another, it has been shown in the field that DNAm clocks are able to predict similar outcomes to one another (Bergsma and Rogaeva 2020; Johnson et al. 2022). Even with these measures having overlaps often as low as 10%, this is still significantly higher than the 2% overlap observed between ELDAR’s key CpGs and the Horvath Pan-Tissue, Horvath Skin-Blood, Hannum, and PhenoAge clocks. Not surprisingly, the majority of the overlapped CpGs were within the age-increasing group of CpGs. This could be a result of there being a number of specific CpGs in an island or window that could be methylated. Therefore, analyses looking at specific CpGs might suffer from exchangeability problems. Yet, larger collections of CpGs are dealing with a large number of trends where many CpGs that sample few phenotype bins, which could also lead to less exchangeability. A limitation of the data used and commonly generated is that the use of targeted DNAm arrays could be leaving out potentially relevant CpGs.

While this underlying principal component based signal was validated in a diverse set of cells, a key limitation is that it remains to be verified in the many additional potential cell types. It is possible that our results may be biased towards fibroblast specific changes given our training datasets. However, some of our strongest validation was observed in brain samples. Another key caveat requiring further investigation, is the consistency of the underlying aging signal across the biological sexes. While female samples were included in the primary skin data and many of the validation datasets, the majority of the cultured cells used in training were BJ fibroblast derivatives, making the training cohort primarily male. Finally, given the small sample sizes in most of our datasets, we employed linear methods for identifying latent patterns. However, given the complexity of the aging process, it may be that nonlinear approaches are needed to elucidate key methylation changes in aging, reprogramming, passaging, and/or transformation. It would not be surprising if PC5’s complex reprogramming patterns are the result of a linear approximation of an underlying nonlinear aging phenotype.

In conclusion, this study has shown that an unsupervised approach to DNA methylation analysis can identify an aging signal that can partially disentangle potentially age-related biological phenomena.. This is the first score that we know of trained across a diverse set of cell states without bias towards a singular “ground truth” metric. As a result, we were able to distinguish two aging signatures. One, which (similar to previous measures) increases with aging and decreases with reprogramming. However, we also identified an orthogonal signature of aging that is not directly inverted by reprogramming, suggesting that aging and reprogramming are not mirrored processes. Moving forward it will be critical to incorporate or align additional omics trajectories with age, along with the DNA methylation signature, to generate a more complete understanding of the underlying aging signal (Vandereyken et al. 2023). This will also be key in determining whether the fundamental drivers of aging are part of DNA methylation, or if DNA methylation is the readout from the drivers.

## Materials and Methods

### 1. Cells and Cell Culture Conditions

BJ Fibroblasts (CRL-2522) were obtained from ATCC (American Type Culture Collection, USA). Cells were routinely cultured in Dulbecco’s modified Eagle’s medium (DMEM) supplemented with 10% FBS (REF: 10437-028), 1x Gibco MEM Non-Essential Amino Acids (REF: 11140-050), and 1x GlutaMax (REF: 35050079). Once 0.22micron filtered the DMEM, FBS, NEAA, and GlutaMax mix was called complete media. Immortalized BJ fibroblasts were generated from the first passage of BJ fibroblasts using the hTert Cell Immortalization Kit from ALSTEM Cell Advancements (cat: CILV02). All cells were grown at 37°C in humidified incubators with 5% CO2 at either hypoxic (3% oxygen), physoxic (5% oxygen), or normoxic (21% oxygen) levels depending on the cell line. Cells would have media exchanged every 3 days until the cells achieved a 90% or greater confluence as determined by imaging with an Olympus EP50 Camera, after which the cells would be passaged. For subculturing the cells would be treated with TryplE Express (REF: 12604-021) at 1ml per 25cm^2 for 6 minutes to lift them off the flask, then equal (double) volume complete media would be added to quench the TryplE. BJ fibroblast sets 1 and 2 came from the ATCC in different lots 2 years apart. The cell suspension would then be centrifuged at 300 RCF for 5 minutes. A DeNovix cell counter was used with Trypan Blue to determine the cell viability and count between subcultures, this was logged to calculate the PDL. The population doubling level (PDL) of the cells was determined by the PDL Equation: PDL = 3.32(log((total live cells collected)/(cells seeded))) + (original seed of line OR prev. PDL). This was annotated as the cumulative population doubling (cPD) in the figures. A caveat to this measure is that for senescent cells which would otherwise have a constant doubling level, their cPD is an estimation taken by determining the average amount of doublings the cell line had per day prior to senescence, and dividing by the days the cells had been in tissue culture. Between subculture events excess cells would be suspended in CryoStor freezing solution (REF: 210502) and frozen in aliquots of 500,000 cells in 1mL CryoStor. Mycoplasma testing was conducted upon receipt of a new cell line and after every month of subculture.

### 2. Cell Lysis and DNA Extraction

DNA was isolated from BJ fibroblasts with a DNeasy Blood and Tissue Kit (Qiagen 69504), according to the manufacturer’s instructions or by MagMAX DNA Multi-Sample Ultra 2.0 Kit (Thermo Fisher Scientific) on the KingFisher (Thermo Fisher Scientific) instrument using the MMX_Ultra2_Cell_Tissue_96_Flex protocol. The DNA was then quantified either by Qubit (Thermo Fisher Scientific) or Quant-it dsDNA-BR Assay (Thermo Fisher Scientific). The DNA was then quality checked by either NanoDrop (Thermo Fisher Scientific) or TapeStation (Agilent). All samples were assigned a code before being randomized for sequencing to control for batch effects.

### 3. Methylation profiling

The DNA samples were then adjusted to a concentration of 25ng/uL at 20uL final volume before submission to the genomics hub for processing on either the EPIC array or Mammalian array for DNA methylation sequencing. There the sample would be bisulfite converted using the EZ-96 DNA Methylation-Lightning™ MagPrep (Zymo). This is followed by the amplification of the converted DNA, fragmentation, precipitation, resuspension, and hybridization to the chip. The bead chips are then run on an iScan or similar Illumina sequencing device.

### 4. Data Processing

The DNA methylation data from the EPIC array was processed using the SESAME (SEnsible Step-wise Analysis of DNA MEthylation) computational tool(Zhou et al. 2018). SESAME was used for preprocessing of raw sequencing reads, alignment, quality control, and identification of differentially methylated regions. Processing was done in RStudio version 4.2.2. The methylCIPHER package was used to calculate the DNA methyl age of the cells according to existing clocks in the literature (Thrush et al. 2022).

### 5. Data Analysis

The ’prcomp’ function in R was used to perform Principal Components Analysis (PCA) on the methylation data. PCA is a statistical procedure that uses an orthogonal transformation to convert a set of observations of possibly correlated variables into a set of values of linearly uncorrelated variables called principal components (Jolliffe, 2017). This technique was used to emphasize variation and bring out patterns with the methylation. The transformation leads to the creation of a two matrix decomposition, where the CpG loadings are obtained from the right singular matrix, and the PC scores are obtained via the projection of the original methylation data onto the right singular matrix.

### 6. CpG and Chromatin State Enrichment

The top 1000 loading CpGs from the PCs of interest were annotated using ChromHMM (Ernst and Kellis 2017). The percent composition of CpG location and Chrom state within the top subset were compared to proportion within the whole dataset to determine the location and states that were enriched within the top loading CpGs.

### 7. Data Accessibility and Usage

All data utilized in the above work are summarized as follows: training: Serial passaged BJ fibroblasts [this study: GSE268145, n = 10 samples], primary sun-exposed dermal samples [GSE52980, n = 20], iPSC time course [GSE54848, n = 18], HRAS cohort samples [GSE91069, n = 15]; validation: Serially passaged BJ fibroblasts [this study: GSE268145, n = 44], serially passaged H-Tert immortalized BJ fibroblasts [this study: GSE268145, n = 22], neuronal differentiation time course [GSE158089, n = 14], serially passaged astrocytes [GSE226079, n = 29], serially passaged H-Tert immortalized astrocytes [GSE226079, n = 45], iPSC time course [GSE54848, n = 30], Sendai mediated reprogramming time course [GSE165180, n = 22], transient OSKM (4F) expression time course [GSE165180, n = 96], liver samples [GSE48325, n = 85], developing brain samples, brain samples <1 in age, [GSE74193, n = 100], brain samples [GSE74193, n = 575], blood samples [GSE40279, n = 656], sun-protected dermal samples [GSE52980, n =20], sun-exposed epidermis samples [GSE52980, n = 19], and sun-protected epidermis samples [GSE52980, n = 19].

## Supporting information

High/Low Loading CpGs List

## List of Abbreviations

ELDAR: Epigenetic Latent signals of Differentiation, Aging, and Reprogramming
PCA: Principal component analysis
PC: Principal component
PDL: Population doubling level
cPD: cumulative population doubling
MEF: Mouse embryonic fibroblast
4F: OSKM
OSKM: Oct4, Sox2, Klf4, and C-Myc
Trans4F: OSKM expression transiently based on Dox induction
DNAm: DNA methylation
Dox: doxycycline
DMEM: Dulbecco’s modified Eagle’s medium
NEAA: Non-Essential Amino Acids
FBS: Fetal bovine serum

## Declarations

All authors are employees of Altos Labs.

## Authors’ Contributions

Conceptualization: P.N. and M.E.L. Methodology: P.N., M.E.L., K.T.-E., and V.G. Investigation: P.N., V.G., and M.E.L. Visualization: P.N. and M.E.L. Funding acquisition: M.E.L. Project administration: M.E.L. Supervision: M.E.L. Writing—original draft: P.N. Writing—review and editing: P.N., M.E.L., K.T.-E., and V.G.

## Acknowledgements

Contribution is based on authorship. We thank our academic and commercial partners for access to data for validating our study.

---

**Supplementary Figure 1:**
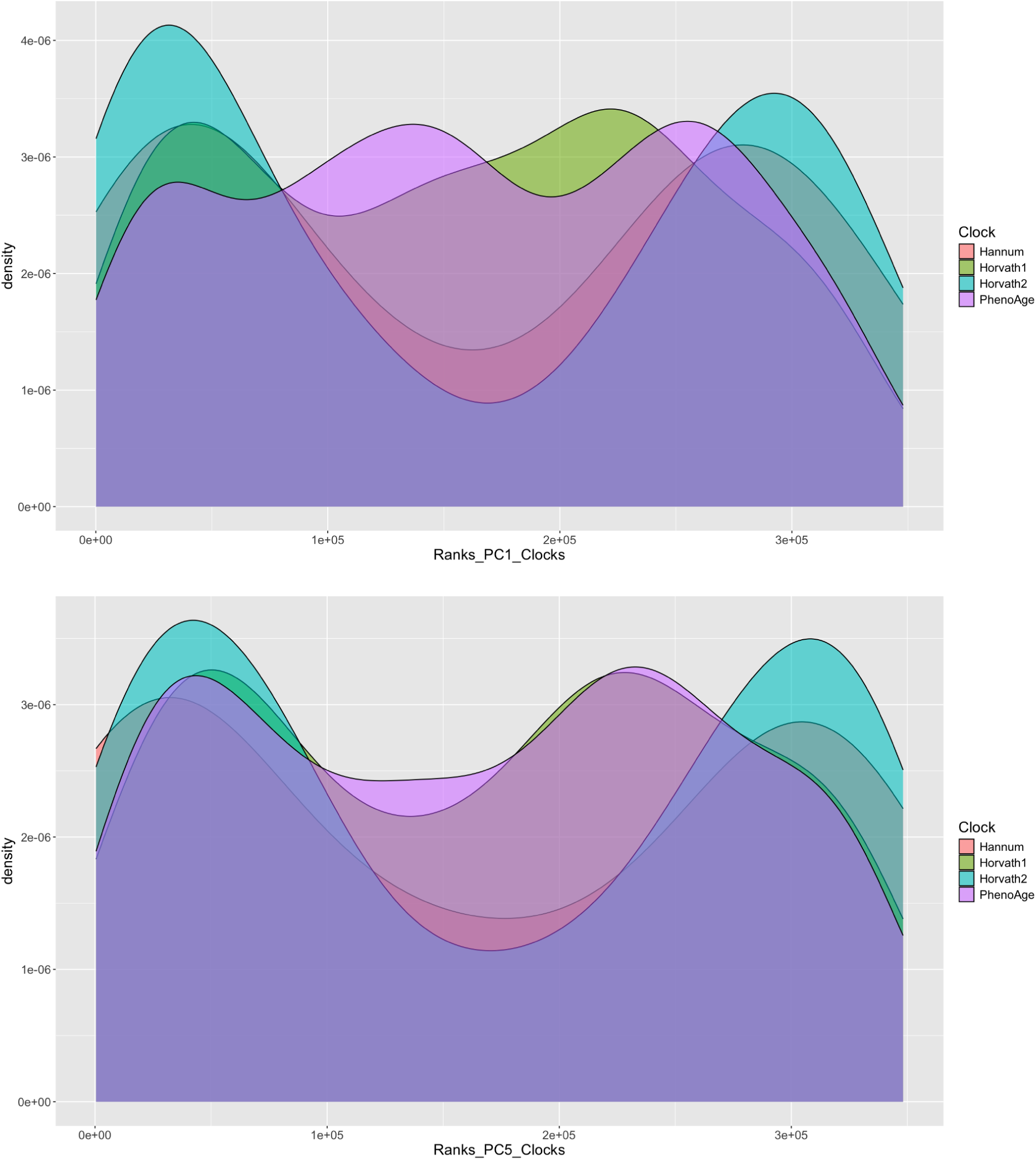
Density of DNAm clock CpGs in ELDAR’s loadings The Horvath2 and Hannum clocks have a correlation with the PC1 and PC5 measures presented here, while there is a negligible correlation with the Horvath1 and PhenoAge clocks. This was determined by arranging all the CpGs in the data by their PC loading and comparing them to the clock CpGs. This supports the overlap found between the arbitrary cutoff of the top and bottom 1000 CpGs with the Hannum, Horvath Skin-Blood, Horvath Pan-Tissue, and PhenoAge clocks.

